# *De novo* ORFs are more likely to shrink than to elongate during neutral evolution

**DOI:** 10.1101/2024.02.12.579890

**Authors:** Marie Kristin Lebherz, Bharat Ravi Iyengar, Erich Bornberg-Bauer

**Affiliations:** Institute for Evolution and Biodiversity, University of Münster, Hüfferstrasse 1, 48149 Münster, Germany; Department of Protein Evolution, Max Planck Institute for Biology Tübingen, Max-Planck-Ring 5, 72076 Tübingen, Germany

**Author notes:** Equal contribution (Marie Kristin Lebherz, Bharat Ravi Iyengar).

**Keywords:** *De novo* gene emergence, gene evolution, protein evolution, mathematical modeling, genomics

## Abstract

For protein coding genes to emerge *de novo* from a non-genic DNA, the DNA sequence must gain an open reading frame (ORF) and the ability to be transcribed. The newborn *de novo* gene can further evolve to accumulate changes in its sequence. Consequently, it can also elongate or shrink with time. Existing literature shows that older *de novo* genes have longer ORF, but it is not clear if they elongated with time or remained of the same length since their inception. To address this question we developed mathematical model of ORF elongation as a Markov-jump process, and show that ORFs tend to keep their length in short evolutionary timescales. We also show that if change occurs it is likely to be a truncation. Our genomics and transcriptomics data analyses of seven *Drosophila melanogaster* populations is also in agreement with the model’s prediction. We conclude that selection could facilitate ORF length extension that may explain why longer ORFs were observed in old *de novo* genes in studies analysing longer evolutionary time scales.

**Significance:** New protein coding genes can emerge from non-genic DNA through a process called *de novo* gene emergence. Genes thus emerged usually have a small open reading frame (ORF). However, studies show that *de novo* genes with an older evolutionary origin have longer ORF than younger genes. To understand how ORF length evolves, we use a combination of mathematical modeling and population level genome data analysis. We find that in the absence of evolutionary selection, ORFs tend to become shorter than becoming longer. Therefore, long ORFs are probably selected by evolution to be retained in the genome.

## Introduction

*De novo* gene birth is phenomenon by which new protein coding genes can emerge from previously non-genic regions of the genome (Van Oss and Carvunis, 2019; Schmitz and Bornberg-Bauer, 2017). This process, that was long thought to be unlikely (Jacob, 1977), is now being increasingly better documented (Carvunis *et al*., 2012; Zhao *et al*., 2014; Neme and Tautz, 2013, 2016; Vakirlis *et al*., 2017; Gubala *et al*., 2017; Baalsrud *et al*., 2017; Prabh and Rödelsperger, 2019; Witt *et al*., 2019; Vakirlis *et al*., 2020; Lange *et al*., 2021; Blevins *et al*., 2021; Wacholder *et al*., 2023). For any stretch of DNA to qualify as a putative protein coding gene, it needs to be transcribed, as well as contain an open reading frame (ORF). Further, for a gene to be considered a *bona fide* protein coding gene, its ORF also needs to be translated. While protein coding genes that arise from existing protein coding genes can inherit the sequence features necessary for transcription and translation from their ancestors, a prospective *de novo* gene must evolve them sequentially through random mutations.

A novel protein coding gene can be fixed in a species, if its products increase the fitness of the organism. Like the features required for transcription and translation, a *de novo* gene cannot inherit the sequence features responsible for fitness effect from an ancestral gene. Although some *de novo* genes have been known to increase the fertility or survivability of an organism (Gubala *et al*., 2017; Baalsrud *et al*., 2017; Lange *et al*., 2021), little is known about the fitness effects of young *de novo* genes. While a tiny fraction of young *de novo* genes may be beneficial to the organism by chance, most of them may not affect organismal fitness at all, or can even be detrimental by causing proteotoxicity (Bucciantini *et al*., 2002; Hartl, 2017).

A protein’s activity is closely linked to how it folds. An established study on protein folding suggests that globular proteins are most likely to fold if they contain more than 70 but less than 2000 residues, and that there exists an optimal protein length that is the best for folding (Dill, 1985). While our understanding of protein folding has advanced significantly, it is still not clear how protein length affects its folding. Unfolded proteins can misfold and cause proteotoxicity (Bucciantini *et al*., 2002; Hartl, 2017). However, not all proteins need to be well folded to be functional (Dyson and Wright, 2005; Wright and Dyson, 2014). For example, many structurally disordered proteins are involved in cell signaling (Wright and Dyson, 2014).

Several studies have shown that conserved proteins are typically longer than putative *de novo* proteins. Furthermore, evolutionarily older *de novo* genes encode longer proteins than younger genes (Carvunis *et al*., 2012; Neme and Tautz, 2013; Zhao *et al*., 2014; Vakirlis *et al*., 2017; Dowling *et al*., 2020; Heames *et al*., 2020; Blevins *et al*., 2021; Middendorf and Eicholt, 2024). It is relatively easy to understand why the protein coding region (ORF) of young genes is short. That is because the likelihood of finding an ORF by chance, as well as the likelihood of ORF emergence, reduces exponentially with ORF length (Iyengar and Bornberg-Bauer, 2023). Thus it is not known whether ORFs of older genes become longer with age, or if only the ones with long ORFs, however less likely they may be in the first place, are the ones that are ultimately fixed.

We attempt to address this question in this study. To this end, we use a combination of mathematical modeling and analysis of genome sequencing data. Mathematical models are useful to understand processes that cannot be easily explained through intuition alone. For example, in a recent work we showed that during *de novo* gene emergence, the likelihood of transcription emerging before an ORF, is higher than *vice versa* (Iyengar and Bornberg-Bauer, 2023). In this study, we developed a mathematical model of ORF length change in *de novo* genes. Because little is known about how *de novo* genes affect organismal fitness, our model is based on the assumption of evolutionary neutrality. Although this does not explain evolutionary dynamics of *de novo* genes through selection, it does provide a good null hypothesis against which observations could be tested. To validate some of our model’s predictions we analyse the genome data for seven *D. melanogaster* populations, and identify how ORF length changes in a short evolutionary timescale. These datasets were generated in a previous study that created inbred lines from a sample of *D. melanogaster* populations from seven different geographical locations, sequenced their genome and transcriptome using deep sequencing, and identified several putative *de novo* protein coding genes in their genomes (Grandchamp *et al*., 2023a). Our analyses of populations, instead of species, allows us to study young *de novo* genes that may not yet be subjected to selection.

Using our two-way approach, we found that ORF of young *de novo* genes are more likely to become shorter than becoming longer. This suggests that neutral evolutionary theory alone cannot explain why older *de novo* genes have longer ORF, and thus selection must be considered to explain this outcome.

## Results

### Development of the mathematical model

We developed a mathematical model to study the dynamics of ORF length evolution, under the assumption that no evolutionary selection occurs for any ORF (neutrality). Specifically, we modelled a Markov jump process, where an ORF can become longer or shorter, or remain of the same length, at any discrete generation. The likelihood of ORF length change is described by a “transition probability”, which is a function of initial ORF length and final ORF length. ORF length can change via mutations, transposition, and even chromosomal recombination. In this study, we focus on ORF length changes that arise due to mutations, and more specifically nucleotide substitutions that are the most frequent kind of mutations. The likelihood of different nucleotide substitutions in a specific genomic locus, depends on mutation rate bias as well as nucleotide composition of the locus (Iyengar and Bornberg-Bauer, 2023). Therefore the ORF length transition probabilities also depend on these two parameters. We used mutation rate bias data from two different organisms – the budding yeast, *Saccharomyces cerevisiae* and the fruitfly, *Drosophila melanogaster* for calculating transition probabilities. We denoted nucleotide composition as GC-content or the frequency of DNA trimers in the intergenic regions from each of the two organisms. Although the ORF length transition probability is determined by nucleotide composition and biased mutation rate, it remains constant over time for any one specific locus. Therefore, the probability distribution of ORF length at a specific locus at any one generation depends only on the same distribution at the previous generation (hence length change is a Markov process).

An ORF can be extended or truncated from both ends through gain or loss of start and stop codons (Figure 1). For example, an ORF extends from the 3’ end if it loses its stop codon, there exists another in-frame stop codon in its 3’ untranslated region (UTR), and no other stop codon exists between the old and the new stop codon positions. For the ORF to shorten from the 3’ end it only needs to gain a premature stop codon. The mechanisms of ORF extensions and truncations from the 5’ end are more diverse than that from the 3’ end. For example, a 5’ extension can occur if a new in-frame start codon emerges in the 5’UTR and there exist no stop codons between the new and the old start codons. A 5’ extension could also occur if an ORF fuses with an in-frame upstream ORF. This can happen if the stop codon of the upstream ORF is lost and there are no stop codons in the intervening sequence between the two ORFs. Finally, a 5’ extension can also occur if the RNA itself is extended from the 5’ (for example, through an alternative transcription start site), that gives rise to a new in-frame start codon in the RNA. Conversely, 5’ truncations can occur via loss of start codon either due to mutations or due to RNA truncation, and splitting of an ORF into two shorter ORFs by a gain of stop codons between two start codons. We note that an ORF can also fuse with a downstream ORF after loss of stop codon, but this mechanism is in principle identical to that of ORF extension to the next in-frame stop codon in an untranslated region. Finally, ORF length can also change due to alternative splicing.

**Figure 1:**
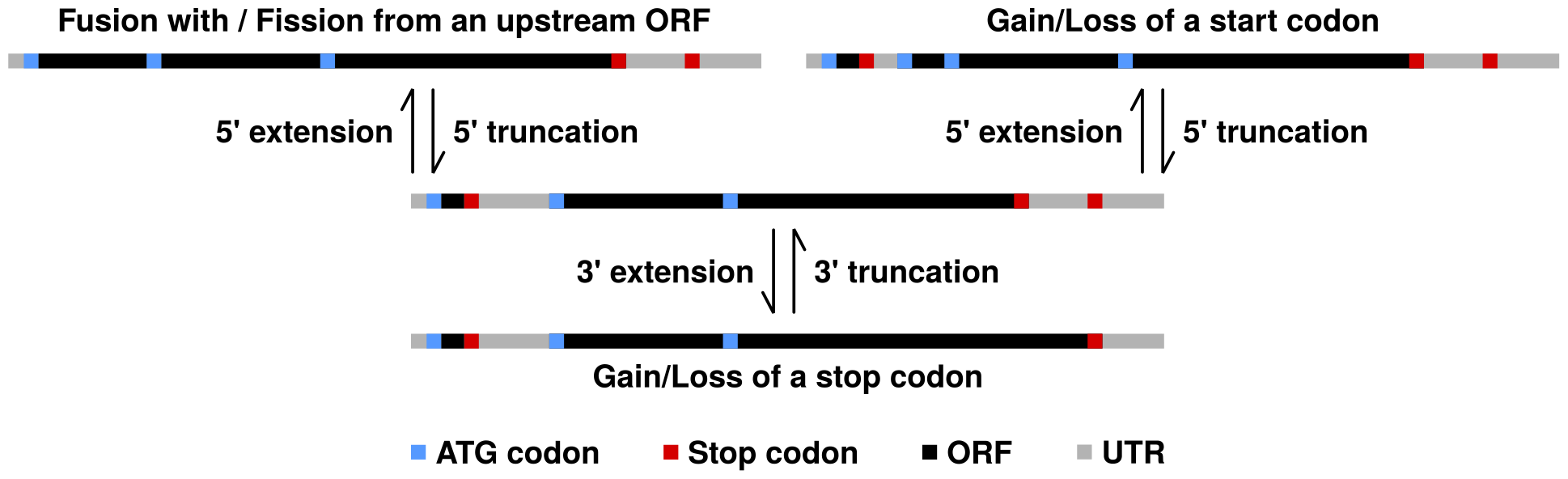
Schematic of the differnet mechanisms of ORF length change, that result due to gain and loss of start and stop codons. We have depicted ATG (start) codons, stop codons, ORF body and UTRs, using blue, red, black and grey colors, respectively.

We modelled the different mechanisms of ORF length change that result due to mutations in start and stop codons (that is, excluding alternative transcription start/end and splicing). To this end we used gain, loss and stationary probabilities of start and stop codons (Iyengar and Bornberg-Bauer, 2023). We analysed the length change of ORFs in the range with a minimum of 3 codons (theoretical minimum), and a maximum of 900 codons, which is an unusually large length for *de novo* ORFs. We defined length transition probabilities within this range. Specifically, we defined a transition matrix (*M*) where the rows and the columns denote ORF length and the elements (*M*_*ij*_, Equation 7) denote the transition probability (*i → j*). Based on the properties of Markov processes, the length transitions over multiple generations (*n*) can be described by the *n*^*th*^ power of the transition matrix (*M* ^*n*^).

### ORFs tend to become shorter with time than to become longer

We used our Markov model to understand how the length of an ORF changes with time, given the ORF already exists. Specifically using the transition matrix (*M*), we calculated the probability that at a future state *F*, an ORF remains of the same length (*F* ^0^), becomes longer (*F* ^+^, to any larger length) or becomes shorter (*F*^*−*^, to any smaller length). These probabilities that depend on the initial ORF length (*i*), and the number of generations (*n*), are descibed as:

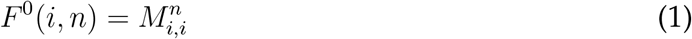

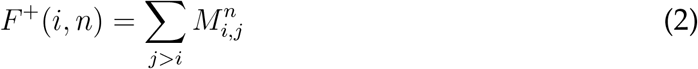

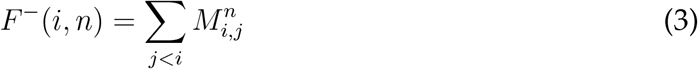

We found that the probability that ORF length changes in one generation is several orders of magnitude (10^6^ – 10^8^) smaller than the probability that it does not. This is understandable because mutations are rare in most organisms (less than 1 mutation in 10^8^ base pairs of DNA per generation; Schrider *et al*., 2013; Zhu *et al*., 2014; Jee *et al*., 2016). Accordingly, we found that for all ORFs, irrespective of their initial length and nucleotide composition, their length tends to remain constant even after several generations (data not displayed). This was the case for our probability estimates using the parameters from both the organisms – *D. melanogaster* and *S*.*cerevisiae*. Next, we investigated when the length does change then whether it increases or decreases. We found that any ORF containing at least 28 codons is more likely to be truncated than extended (Figure 2). This minimum ORF length is a function of both the number of generations and the nucleotide composition. For example, based on our *D. melanogaster* parameters, ORFs present in a locus with 30% GC-content, and containing at least 18 codons are likely to be truncated in one generation. After 2 *×* 10^7^ generations all ORFs with at least 9 codons are likely to be truncated, irrespective of the nucleotide composition (Figure 2A). We found similar trends using our *S. cerevisiae* parameters (Figure 2B). For example, up to *∼* 1.7 *×* 10^6^ generations the shortest ORF present in a locus with 30% GC-content that is more likely to extend than to truncate, contains 16 codons. After 2 *×* 10^7^ generations, all ORFs with at least 9 codons are likely to be truncated, irrespective of the nucleotide composition.

**Figure 2:**
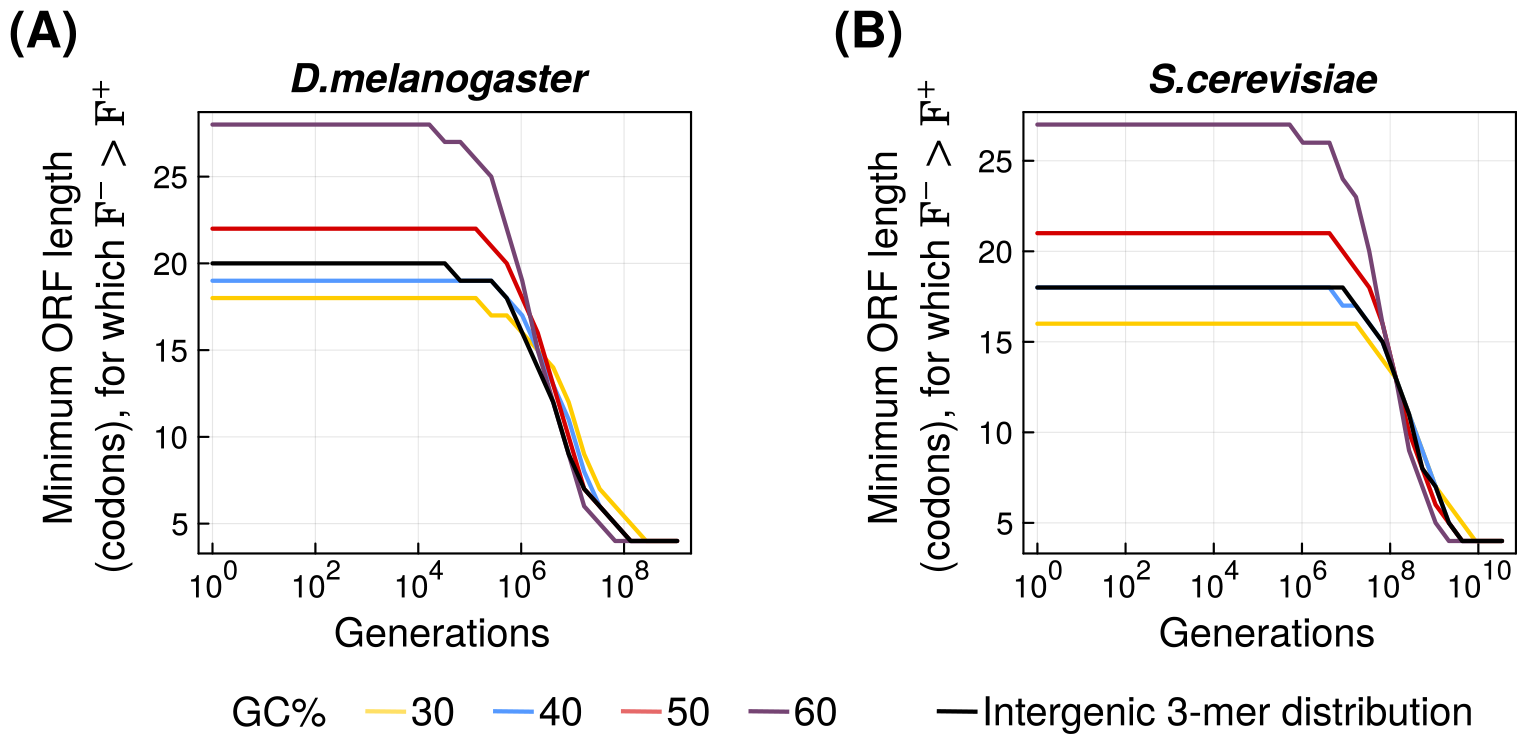
ORFs tend to become shorter with time than to become longer. Vertical axes show the length of the shortest ORF (in codons) that is more likely to become shorter than to elongate (*F*^*−*^ *> F* ^*+*^; Equations 2 & 3). Horizontal axes show the number of simulated generations of **(A)***D. melanogaster* and **(B)** *S. cerevisiae*, in log scale. Line colors denote the nucleotide composition of the locus: yellow = 30%GC, blue = 40%GC, red = 50%GC, purple = 60%GC, and black = intergenic trimer frequencies.

**Figure 3:**
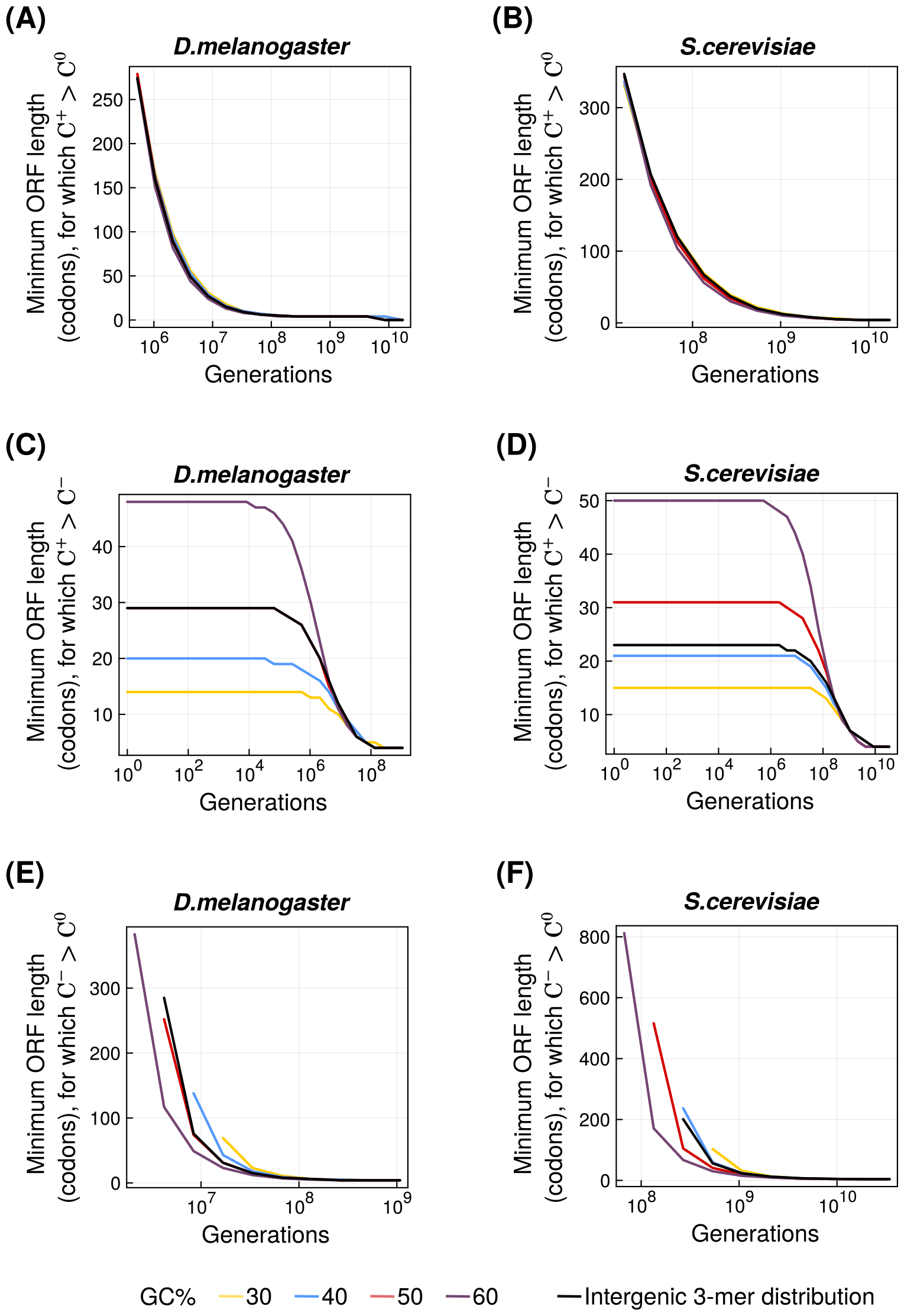
ORFs are more likely to emerge from the extension of smaller ORFs than from truncation of larger ORFs. The shortest ORF (vertical axis) for **(A)** *D. melanogaster* and **(B)** *S*.*cerevisiae* that is more likely to have descended from any shorter ORF than from an ORF of the same length (*C*^*+*^ *> C*^0^). The shortest ORF (vertical axis) for **(C)** *D. melanogaster* and **(D)** *S. cerevisiae* that is more likely to have descended from any shorter ORF than from any longer ORF (*C*^*+*^ *> C*^*−*^). The shortest ORF (vertical axis) for **(E)** *D. melanogaster* and **(F)** *S. cerevisiae* that is more likely to have descended from any longer ORF than from an ORF of the same length (*C*^*−*^ *> C*^0^). See Equations 4 *–*6 for details. Horizontal axes in all panels show the number of simulated generations in log scale. Line colors denote the nucleotide composition of the locus: yellow = 30%GC, blue = 40%GC, red = 50%GC, purple = 60%GC, and black = intergenic trimer frequencies.

### ORFs are likely to be a product of extension than of truncation

We next analysed whether an ORF of a given length originated as an ORF of a different length (shorter or longer) or the same length. To this end, we calculated three probabilities that describe the current state of the ORF (*C*). First, the probability *C*^0^(*i, n*) that an ORF of a given length (*i*) remains of the same length after *n* generations. Second, the probability *C*^+^(*i, n*) that any ORF with length *j < i* extends to an ORF of length *i* in *n* generations. Finally, the probability *C*^*−*^(*i, n*) that any ORF with length *j > i* truncates to an ORF of length *i* in *n* generations. If *P*_*ORF*_(*i*) denotes the probability of finding an ORF containing *i* codons (Iyengar and Bornberg-Bauer, 2023) then the three current state ORF probabilities are described as:

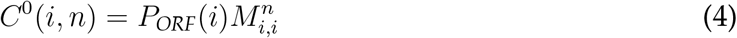

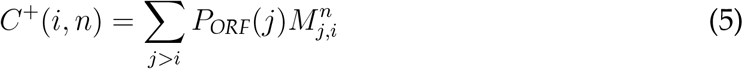

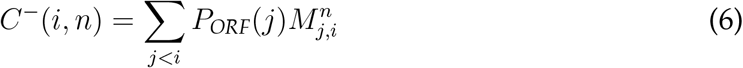

The likelihood of finding an ORF decreases exponentially with its length (Iyengar and Bornberg-Bauer, 2023). Therefore we first asked if it is possible that the scenario where an ORF is extended from smaller ORFs in *n* generations (*C*^+^), is more likely than the scenario where an ORF’s current length is same as what it was *n* generations ago (*C*^+^). From our previous analysis we know that ORF probabilities depend on ORF length. Specifically, longer ORFs are less likely to have emerged from an ORF of the same length. In contrast, longer an ORF is, more the chances are that it emerged from a shorter ancestor. Therefore, we identified the smallest ORF (*i*) for which *C*^+^(*i, n*) (Equation 5) is greater than *C*^0^(*i, n*) (Equation 4), at different generations (*n*). We found that *C*^0^ is greater than *C*^+^ for all the analysed ORF lengths until *∼* 1.4 *×* 10^5^ simulated *D. melanogaster* generations. From this generation onwards the minimum ORF length for which *C*^+^ is greater than *C*^*−*^, decreases with increasing generations. For example, it decreases from 707 codons at *∼* 1.4 *×* 10^5^ generations to 4 codons at *∼* 10^8^ generations, for a locus with a GC-content of 40%. We found that these ORF lengths varied only modestly with varying nucleotide composition. We made similar observations using our parameter estimates from *S. cerevisiae*. However, *C*^+^ overtakes *C*^0^ only after 8.4 *×* 10^6^ simulated *S. cerevisiae* generations. We note that for organisms, *C*^+^ overtakes *C*^0^ when the product of the number of generations and the mutation rate exceeds 0.001.

Next, we asked if it is possible that at least some ORFs are more likely to arise due to truncation of larger ORFs than due to extension of smaller ORFs. To this end, we compared the probabilities *C*^*−*^ and *C*^+^ and calculated the smallest ORF length for which *C*^+^ is greater than *C*^*−*^. We found that only short ORFs are likely to be a product of truncation. For example, after one simulated *D. melanogaster* generation, any ORF present in a locus containing 40% GC and a maximum size of 20 codons is likely to have truncated from larger ORFs. This remains so until *∼* 2.6 *×* 10^5^ generations after which even smaller ORFs are more likely to be products of extension. Our analysis using *S. cerevisiae* parameters also revealed similar findings. Like in case of *D. melanogaster*, an ORF present in a locus with 40% GC and with a minimum length of 21 codons, is more likely to have been extended from smaller ORFs than truncated form larger ORFs. However, this minimum length starts decreasing only at *∼* 10^7^ generations. In both the organisms the likelihood of an ORF being a product of extension relative to that of truncation, reduces with increasing GC-content.

Finally, we asked if the probability of an ORF originating from a larger ORF (*C*^*−*^) can be greater than the probability of it originating from an ORF of the same size (*C*^0^). Although non-intuitive, this scenario is indeed possible if the number of generations is large enough. For example, at *∼* 4.2 *×* 10^6^ simulated *D. melanogaster* generations, any ORF present in a locus with 40% GC and is longer than 137 codons, is most likely to have been truncated from larger ORFs than to have originated form an ORF of the same length. Our analysis with *S. cerevisiae* as a model, also produced similar results. The number of generations where *C*^*−*^ exceeds *C*^0^ is inversely proportional to the mutation rate. We also find that the likelihood of being truncated (*C*^*−*^) relative to being extended (*C*^+^), increases with increasing GC content.

### Length changes in *D. melanogaster de novo* ORFs are more frequent than expected

To understand how ORF length changes in actual organisms, we analysed a recently published dataset on *de novo* transcripts in seven inbred *D. melanogaster* lines obtained from seven geographically distinct populations (Grandchamp *et al*., 2023a,b). These seven lines were predicted to have diverged from a common ancestor ca. 13000 years ago (Grandchamp *et al*., 2023b). This corresponds to approximately 333400 generations (Fernández-Moreno *et al*., 2007). Next, we identified all possible ORFs (*≥*30nt) in the novel transcripts from each of the seven lines, and sorted them into groups of orthologous ORF sequences (orthogroups) based on sequence homology and synteny (that is, identical flanking genes). We also identified untranscribed ORF orthologs by analysing the genomic regions syntenic to those of transcribed orthologs, and included them in the orthogroups. We thus constructed 758 orthogroups of which 48%, had orthologous ORFs from all seven lines (Figure 4A). Next, we analysed the lengths of the ORFs within each orthogroup. Specifically, we identified orthogroups in which all the constituent ORFs had identical lengths, and those that had ORFs with different lengths. Most orthogroups (77%) did not have any length variation between their constituent ORFs (Figure 4A). This observation only qualitiatively agrees with our model, because the percentage of orthogroups with length variation (23%) between their constituent ORFs, was significantly higher than expected (*∼*4.8%; *P <* 10^*−*6^, Monte-Carlo sampling). Possible reasons for frequent ORF length changes could be variability of transcription start sites and splice sites, both of which we did not incorporate in the model.

**Figure 4:**
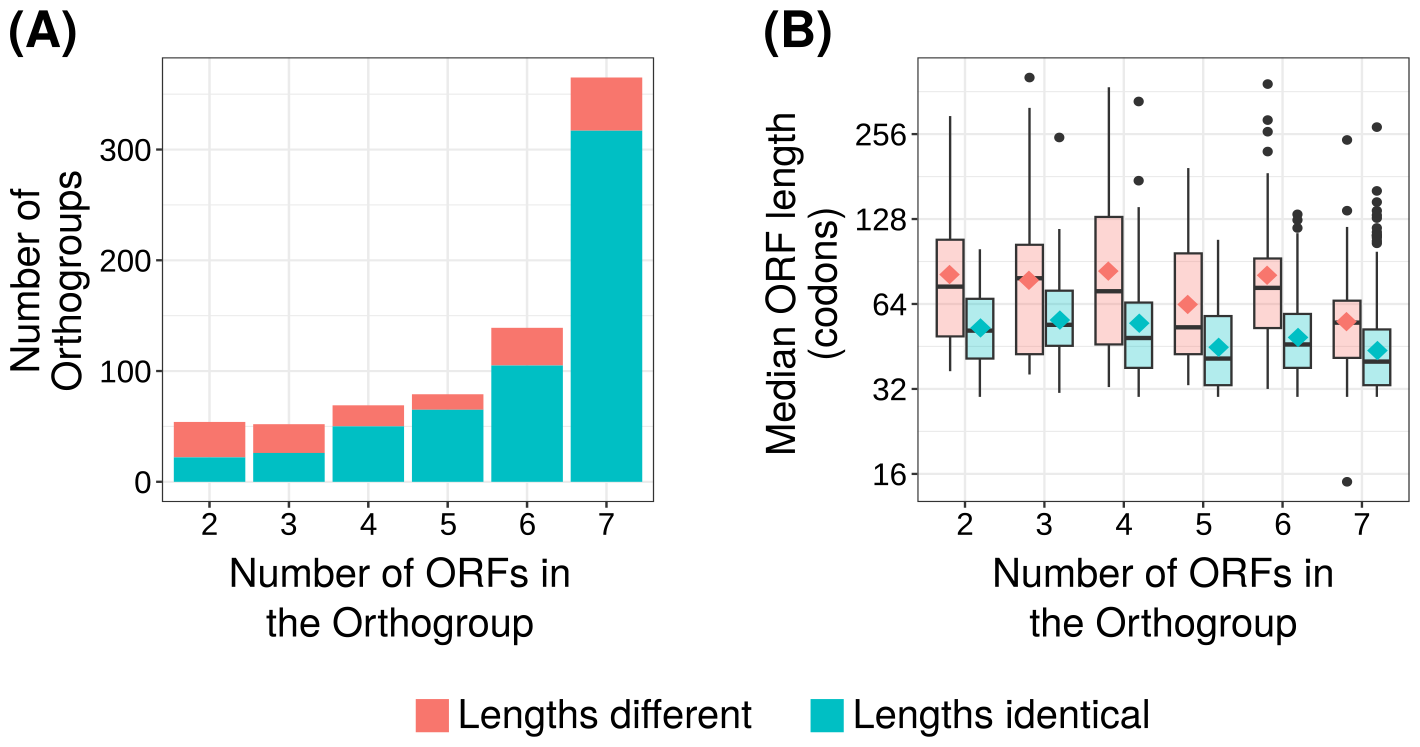
ORFs are more likely to emerge from the extension of smaller ORFs than from truncation of larger ORFs

A few studies analysing *de novo* genes across long evolutionary time scales comprising speciation events, report that *de novo* ORFs become longer with evolutionary age (Carvunis *et al*., 2012; Neme and Tautz, 2013; Zhao *et al*., 2014; Vakirlis *et al*., 2017; Dowling *et al*., 2020; Heames *et al*., 2020; Blevins *et al*., 2021; Middendorf and Eicholt, 2024). In contrast, our model suggests that an ORF is more likely to become shorter than to become longer with age (Figure 2). To further understand whether truncation is more common than extension, we analysed our orthogroups that contain ORFs with different lengths. Orthogroups that contain ORFs from more number of *D. melanogaster* lines may be evolutionarily older than those that contain ORFs from fewer lines. This is especially evident for orthogroups that contain ORFs from all seven lines including the outgroup. Therefore, we analysed the correlation between the number of orthologous ORFs within an orthogroup and the median length of these ORFs. We did so for both the orthgroups with length variation and those without length variation. We found a significant negative correlation between median ORF length and the number of lines harboring an ORF ortholog (Figure 4B; orthogroups with lengh variation: Spearman’s *ρ* = *−*0.183, *P* = 7.8 *×* 10^*−*3^; orthogroups without lengh variation: Spearman’s *ρ* = *−*0.186, *P* = 2.89 *×* 10^*−*6^).

To more precisely understand if ORFs become shorter with time, we estimated the length of the ancestral ORF for each orthogroup that contains ORFs of different lengths. Specifically, we used a dated phylogenetic tree (Grandchamp *et al*., 2023b) to find out the most recent common ancestor that would have harbored an ORF belonging to a specific orthogroup. For example, if an orthogroup contains ORFs from the Swedish line and the Danish line, then the last common ancestor of both these lines must also contain a homologous ORF. We assign this ORF as the ancestor of the orthogroup. We note that we do not perform ancestral sequence reconstruction but simply assume that an ancestral ORF exists and it can have any possible length. Next, we calculate the probability that the ORFs in an orthogroup could have the length they have, given the phylogeny between the populations and that the ancestral ORF has a specific length. Specifically, we calculate the transition probability than an ancestral ORF of a length (*i*) gives rise to an extant ORF of length (*j*), in the number of generations estimated from the length of the evolutionary path that connects the ancestral population and the extant population (that contains the ORF). We perform the same calculation for every other ORF in the orthogroup, while excluding the branches of the tree that have been already counted. The multiplicative product of these different transition probability values indicates the likelihood of an ancestral ORF length, such that the most likely ancestral ORF length will produce the largest value. Using this technique, we predicted the most likely ancestral ORF length for every orthogroup, and in turn, the frequencies of truncations and extensions. We found that 70.5% ORFs had the same length as their ancestor, 25% had been truncated from a longer ancestor, and 4.5% had been extended from a shorter ancestor. We emphasize that these results are based on the assumption of evolutionary neutrality.

Overall, our analyses of *D. melanogaster de novo* ORFs suggest that truncation is more likely than extension.

### Length changes in *D. melanogaster de novo* ORFs are larger when they occur at the 5’ end than from the 3’ end

We next focused on the mechanism of ORF length changes. We could expect 3’ extensions to be smaller in magnitude than 5’ extensions because it is more likely to find one of the three stop codons by chance than to find a start codon. However, unlike the loss of stop codons in 3’ extensions, a gain of a new start codon during 5’ extension does not necessitate the loss of the downstream (old) start codon. Furthermore, the mechanisms and probabilities of truncation are different from that of extension (Figure 2, Equations 2 & 3). Because the differences in the mechanisms of 5’ and 3’ length changes are not trivial, we analysed both our model and the *D. melanogaster* data, to understand which change is more frequent and is more in magnitude.

To this end, we first identified the longest ORF in an orthogroup that has ORFs of different lengths, and aligned it to all the shorter ORFs in the same orthogroup, using protein BLAST. Using the alignment, we determined whether the longest ORF is extended from the 5’ end or the 3’ end (or both), relative to the shorter ORFs. We analysed 229 ORF pairs, out of which 142 pairs shared the same start position (62%, 3’ change), 81 shared the same stop position (35.4%, 5’ change), and 6 shared neither of the two termini (2.6%, changed form both the ends). Next, we compared the extent of length changes from the 5’ and the 3’ ends, and found that 5’ changes (median 48 codons) were larger than the 3’ changes (median 21 codons; one-sided Mann-Whitney U-test, *P* = 4.6 *×* 10^*−*4^; Figure 5A). This is in qualitative agreement with our model’s prediction that also shows that large changes are more likely to occur in the 5’ end than from the 3’ end (Figure 5B). Overall, we found that 3’ changes were more frequent but produced smaller length difference between the ORFs.

**Figure 5:**
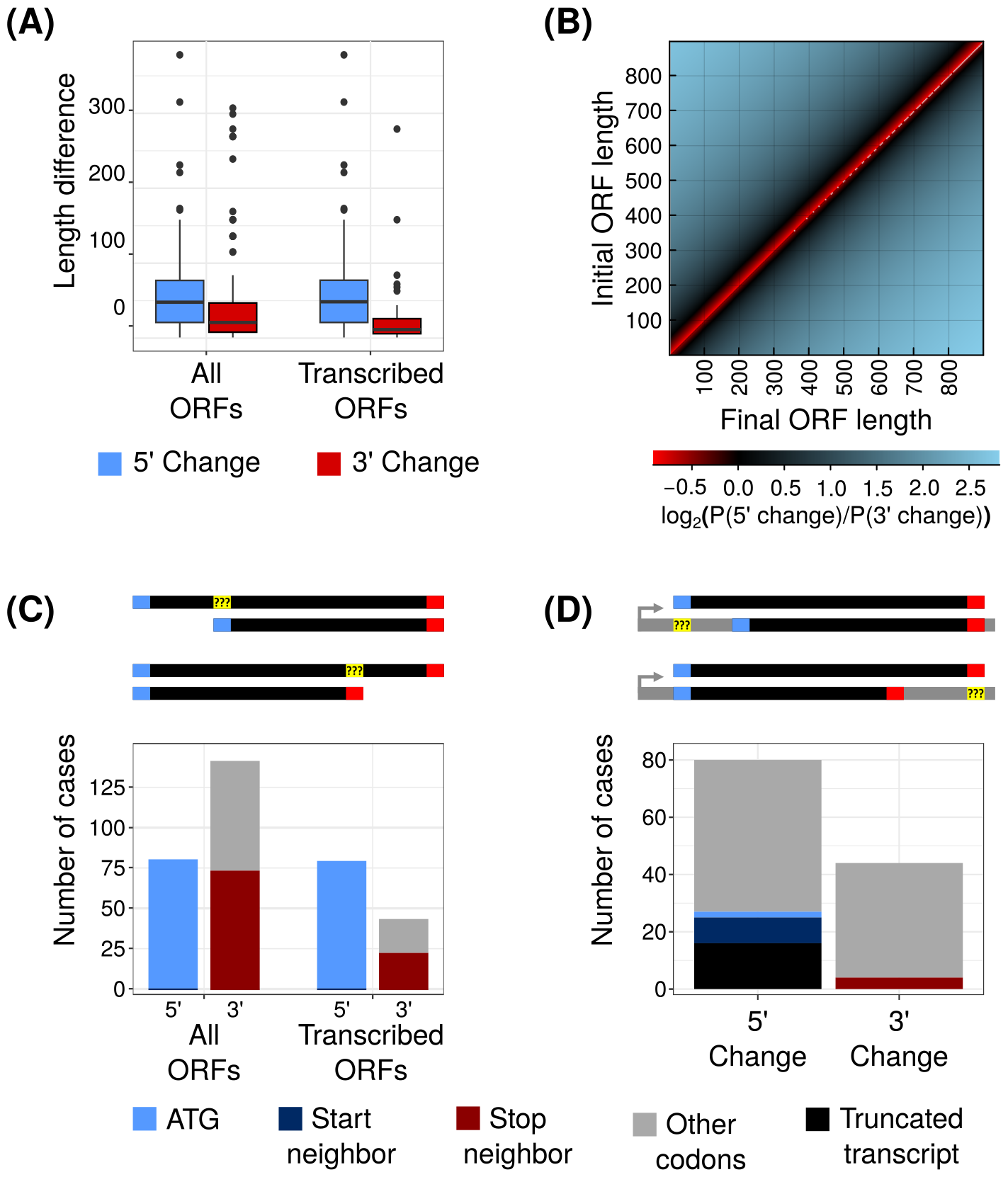
Analyses of ORF length changes from the 5′ end and the 3′ end. **(A)** Boxplots depicting ORF length changes from the 5′ end (blue boxes) are larger (vertical axis than those from the 3′ end (red boxes) in the orthogroup dataset constructed using all ORFs as well as the orthogroup dataset containing only the transcribed ORFs (horizontal axis). The boxes extend from the first to the third quartile, the whiskers have a length equal to 1.5 *×*the interquartile range, the outliers are denoted by black circles, and the black horizontal bar inside each box depicts the corresponding median. **(B)** Heatmap depicting the log_2_ transformed ratio of the probability of a 5′ length change relative to that of a 3′ length change (color scale) for different values of initial (vertical axis) and final (horizontal axis) ORF lengths. Blue shades denote probability of 5′ change being greater than that of 3′ change, while red shades denote *vice versa*. See Equations 8 –9. **(C)** Number of cases in both the datasets (all ORFs and transcribed ORFs, groups in horizontal axis) where the codons of each type in the longest ORF (colors: blue = ATG, dark blue = start neighbor, dark red = stop neighbor, and grey = other codons), map to the terminal codons (5′ or 3′, horizontal axis) of the shorter ORFs. **(D)** Number of cases in the datasets with transcribed ORFs where the codons of each type in the shorter ORFs (same color scheme as **C**) map to the terminal codons (5′ or 3′, horizontal axis) of the longest ORFs. Black bar denotes the number of cases where the position in the short ORF corresponding to the long ORF does not exist in the corresponding transcript.

ORF length could also be altered due to changes in transcription start site. For example, a more upstream transcription start site could cause an inclusion of an in frame start codon, that in turn could extend the ORF from the 5’ end. Because we constructed our orthgroups by first identifying transcribed ORFs, ORF length differences between two orthologous ORFs could exist due to differences in their transcription start sites (TSS). Therefore, we repeated our previous analyses of ORF start and end sites (previous para), with only the transcribed ORFs. We now analysed 130 ORF pairs, and found that 44 were altered from the 3’ end, 80 were altered from the 5’ end and 6 were altered from both the ends. Furthermore, we found that 5’ changes, that are now more frequent, also produced a greater extent of length change (one-sided Mann-Whitney test, *P* = 2.4 *×* 10^*−*6^; Figure 5A)

Next, we analysed the mechanism underlying the ORF length changes. Length changes at the 3’ end could result from gain or loss of a stop codon. To verify if this is the case, we analysed the codon in the longest ORF that overlaps with the stop codons of smaller ORFs in the same orthogroup. Specifically, we asked if this codon is a single nucleotide mutation away from a stop codon (stop neighbor). We performed this analysis with both our full dataset containing all ORFs and the dataset containing only the transcribed ORFs. For both datasets we found that a stop neighbor was present in the longest ORF in *∼*52% of cases (52.1%: all, 52.3%: transcribed only; Figure 5C). Because it is very less likely for a stop codon to mutate into a non-stop codon with more than 1 nucleotide substitutions (*∼* 10^*−*17^), one would expect nearly every codon in the long ORF that overlaps with a stop codon in the short ORF, to be a stop neighbor. Our data analysis suggests that a large proportion of 3’ changes do not occur due to a simple gain or loss of a stop codon. We performed an analogous analysis where we identified the codon in the longest ORF that overlaps with the start codon in the shorter ORFs. We found that in every case (and in both the datasets), the longest ORFs indeed had an ATG (*∼*99%) or a 1nt-neighbor of ATG (start neighbor, *∼*1%) at the position overlapping the start codon in the smaller ORFs (Figure 5C), in accordance with our expectation.

A truncation from the 5’ end could result from either the loss of a start codon (with the availability of a downstream in-frame start codon) or the gain of a stop codon in between two in frame start codons. Furthermore, transcript truncation from the 5’ end (alternative TSS) could also cause ORF truncation. To assess these diverse possibilities, we analysed the dataset containing only the transcribed ORFs, and identified the codon in the 5’UTRs of the short ORFs in an orthogroup that map to the start codon in the longest ORF of the same orthogroup. In the majority of the cases (78 out of total 80: 97.5%) we did not find a start codon at the 5’UTR position overlapping the start codon of the longest ORF, out of which 33.3% were start neighbors, suggesting that a loss of a start codon could have caused the length change (Figure 5D). In 2.5% of the cases the overlapping codon in the 5’ UTR short ORFs was ATG, suggesting that there could have been a gain of stop codon between this ATG and the actual start codon. Interestingly in 20 % of the cases, we found the transcript of the short ORF to be truncated from the 5’ end such that no 5’UTR positions existed that could overlap with the start codon of the longest ORF (Figure 5D). We analysed the 3’ changes analogously and found that in the majority of the cases (40 out of total 44, 90.9%) no stop codon could be identified in the short ORFs in an orthogroup that overlapped with the stop codon of the longest ORF in the same orthogroup (Figure 5D). Possible reasons for this observation could be poor sequence alignment in the 3’UTR, that may result from mechanisms such as alternate splicing.

Overall, our analyses show that ORF length tends to change more from the 5’ end than the 3’ end.

## Discussion

In this study, we aimed to understand how ORF length changes during the course of evolution. To this end, we first developed a mathematical model that predicts ORF length changes under the assumption of evolutionary neutrality (Equation 7). We used the model to ask two questions. First we asked, if given that an ORF of a certain length exists, what the chances are that it retains its length, becomes shorter or becomes longer (Equations 1 – 3). ORFs are more likely to retain their length than to become longer or shorter. This is an expected outcome because mutations are rare in most organisms including the two used in our modeling analyses. However, we found that if ORF length does change then it is more likely to decrease than increase (Figure 2). Our second question pertains to the evolutionary past of the ORF. Specifically, we asked if ORFs originate from ancestral sequences that are shorter, longer, or of the same size (Equations 4 – 6). Our model shows that ORFs are most likely to arise from ancestral ORFs of identical length in short evolutionary timescales. However, the longer in past one goes, the higher the likelihood of an ORF originating from an ORF of dissimilar length becomes. ORFs are especially likely to elongate from shorter ancestors than from longer ancestors (Figure 3). This finding may seem contradictory to that of our first question where we find that ORFs are more likely to become shorter with time. The difference hinges mainly on what we know about an ORF. When we predict the future of the ORF, we are sure about its current length and estimate an average of all possible future outcomes (extensions or truncations). However, when we predict the past states of an ORF, we are unsure about the initial ORF length and assume that they are geometrically distributed such that longer ORFs are exponentially less likely to exist (Iyengar and Bornberg-Bauer, 2023). Thus ORFs are more likely to extend from shorter ancestors.

To validate some of these predictions, we analysed length changes in orthologous *de novo* ORFs from seven *D. melanogaster* lines obtained from seven different populations. We found that most orthogroups contained ORFs of identical length, which is qualitatively in line with our model’s predictions. However, we found more orthogroups with length changes than expected. One possible reason could be that our model ignores mutational events that can change transcription start sites (TSS) and splice sites, that can be important determinants of ORF length change. Indeed, our subsequent analysis shows that in 20% of ORFs with a length change from the 5’, occurs due to differential TSS usage in the corresponding transcripts (Figure 5D). We also analysed the orthogroups to ask if ORFs indeed become truncated with time. To address this question we first used a simple approach where we compared the median ORF length of an orthgroup to the number of ORFs contained in it. We found that orthogroups with more number of ORFs had a smaller median length suggesting that short ORFs are more likely to survive (Figure 4B). Although orthogroups containing ORFs from all seven lines definitely have a relatively ancient ancestor, the orthgroups with fewer ORFs need not necessarily have a more recent ancestor possibly because of widespread ORF loss. Thus widespread survival of short ORFs alone doesn’t suffice to answer if they originated from a longer ancestor. Therefore, we used a more refined approach where we inferred the most likely ancestral length of an orthogroup using our mathematical model and a dated phylogenetic tree of the seven *D. melanogaster* populations (Grandchamp *et al*., 2023b). Using this approach, we quantified truncations that are more likely than extensions.

Our findings stand in contrast to those of several studies that report that ORFs of older *de novo* protein coding genes are longer (Carvunis *et al*., 2012; Neme and Tautz, 2013; Zhao *et al*., 2014; Vakirlis *et al*., 2017; Dowling *et al*., 2020; Heames *et al*., 2020; Blevins *et al*., 2021; Middendorf and Eicholt, 2024). Our model is based on the assumption of neutral evolution, and the *de novo* ORFs we analysed are relatively very young (most of them not even fixed in the species). In contrast, the above mentioned studies consider single genomes from many species that have diverged several millions of years ago. Therefore, evolutionary selection is a very likely explanation for the larger length of old *de novo* ORFs found in these studies, that our analyses do not take into account. Some *de novo* genes may become fixed in a species (or a clade) due to positive selection. Indeed, some *de novo* genes have been experimentally shown to increase organismal fitness (Gubala *et al*., 2017; Baalsrud *et al*., 2017; Lange *et al*., 2021; Wacholder *et al*., 2023). Purifying selection on these genes may prevent the truncation of the ORF. On the other hand, ORF extension may increase the beneficial fitness effects of a gene which can lead to its fixation. For example, C-terminal (3’) extensions that are structurally disordered could protect proteins from deleterious effects of stop codon readthrough (Kleppe and Bornberg-Bauer, 2018). An evolutionary stable gene, especially one that produces a thermodynamically stable protein, can also facilitate ORF extension through a purely neutral evolution. This is possible because protein stability can facilitate evolutionary innovation (Bornberg-Bauer and Chan, 1999; Bloom *et al*., 2006; Zheng *et al*., 2020).

Our study, like every other study, is based on certain approximations and is limited by availability of data. Therefore, one must consider these limitations before applying the findings to a larger scale, and for making broad generalizations. For example, we use GC-content and trimer frequencies to approximate nucleotide distributions. Both these parameters may vary significantly throughout the genome. However, using locus specific nucleotide distributions can predict evolution of the corresponding loci more realistically. Our model also does not take into account all the possible mechanisms that lead to ORF length change. For example, we do not incorporate changes in transcription start site (TSS), primarily because we do not have enough data to model the evolution of TSS. When such data become available the model can be updated. Our data analysis is also limited by the amount of available information. For example, we cannot estimate the exact divergence time between the *D. melanogaster* populations. Moreover, the generation time is unlikely to be similar between populations living in vastly different climatic regions. Furthermore, the populations may not be strictly geographically isolated, as suggested by small genetic variation (F_ST_) between the European populations (Kapun *et al*., 2020).

Despite many simplifications in our analyses, our study provides many qualitative answers, and also a formal basis to test hypotheses through more focused experiments. It also opens up several questions that may be a topic of future research. For example, it may be worth investigating if terminal extensions can indeed improve the fitness effect of a protein or if they are mostly deleterious or neutral. Ultimately, studying the ancestry of different extant *de novo* ORFs could help understand if they were extended from smaller ancestors or if they were born with the same length, and could help dissect the different mechanisms that lead to fixation of *de novo* genes with long ORFs.

## Methods

### Calculation of transition probability

We calculated the probability of ORF length change due to gain and loss of stop codons (Figure 1), in the form of a transition matrix (*M*) whose rows denote the initial ORF length and the columns denote the final ORF length. We generated transition matrices for two organisms – *Drosophila melanogaster* and *Saccharomyces cerevisiae*. For each organism we calculated the transition probabilities using the values of biased mutation rate (*D. melanogaster*: Schrider *et al*.,2013; *S. cerevisiae* Zhu *et al*.,2014), and nucleotide compostion described by four different values of GC-content (30%, 40%, 50% and 60%), as well as the distribution of DNA trimers in the intergenic regions of the organism. We calculated the values of the elements of the transition matrix (*M*_*ij*_, Equation 7) using probabilities of finding, gaining and losing start and stop codons (Table 1; Iyengar and Bornberg-Bauer, 2023).

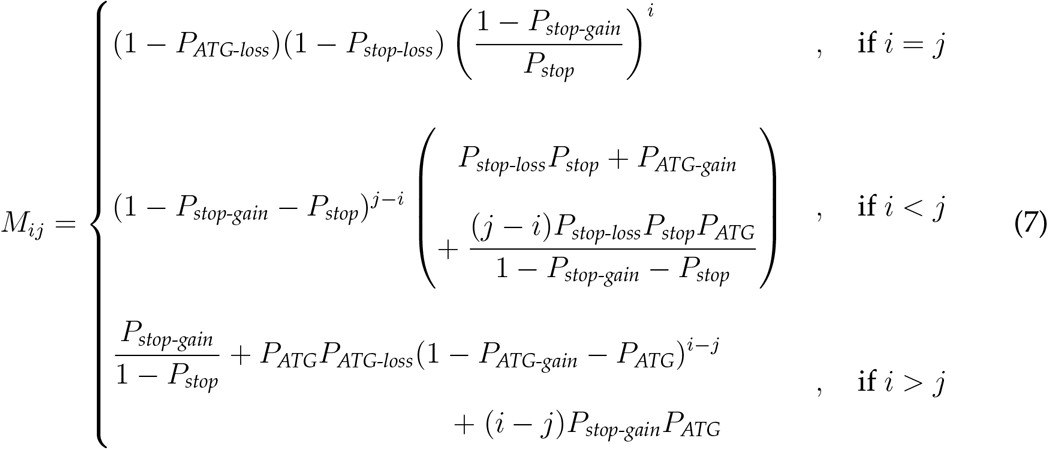

By excluding some terms from Equation 7, we calculated the probability of length transitions from only the 5’ end (Equation 8) or only the 3’ end (Equation 9).

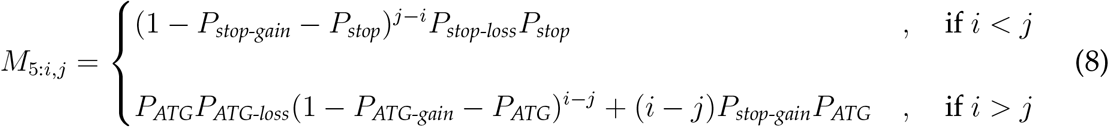

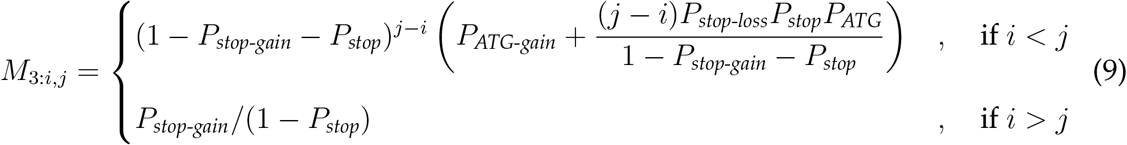

**Table 1:**
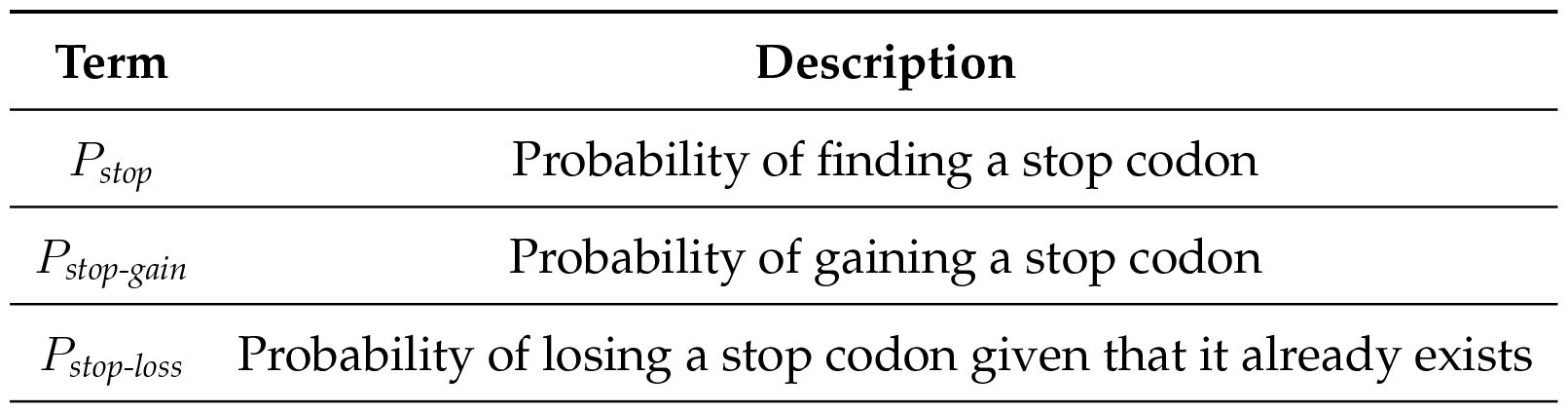
Description of the probability terms used in Equation 7. Here we describe the probabilities associated with stop codons (based on Iyengar and Bornberg-Bauer, 2023). Analogous probability terms for a start codon are denoted by the subscript, *ATG* (instead of *stop*).

| Term | Description |
| --- | --- |
| $P_{stop}$ | Probability of finding a stop codon |
| $P_{stop-gain}$ | Probability of gaining a stop codon |
| $P_{stop-loss}$ | Probability of losing a stop codon given that it already exists |

### Construction of ORF orthogroups containing transcribed ORFs

We identified ORFs in the novel intergenic transcripts originating in seven *D. melanogaster* inbred lines obtained from seven different geographically distinct populations (Lebherz *et al*., 2023). Specifically, we identified all ORFs starting with the canonical start codon (ATG), and containing a total of at least 10 codons (30nt), using the program *getorf* (Rice *et al*., 2000).

Next, we grouped these ORFs into orthogroups. To this end, we used protein BLAST (Altschul *et al*., 1990; Camacho *et al*., 2009) to align ORFs from all the seven lines with each other, at 100% query coverage, 90% sequence identity and an E-value cutoff of 10^*−*4^. We grouped all the BLAST hits into orthogroups such that every ORF in each orthogroup aligned with at least one another ORF in the same orthogroup (based on our hit cutoffs). We discarded orthogroups that contained more than one ORF per line. Next, we analysed the genes flanking the ORF of an orthogroup (synteny), using genome annotations for the seven lines (Grandchamp *et al*., 2023a). If the neighboring genes were identical for all ORFs, we kept them in the same orthogroup, and split them into multiple orthogroups, otherwise. We applied this method to every orthogroup. We further analysed only the orthogroups that contained at least two ORFs.

### Identification of non transcribed homologous ORFs

Because our study focuses primarily on ORF length extension but not gain and loss of transcription, we included non-transcribed ORFs in our orthogroups, that were homologous and syntenic to the transcribed ORFs the same orthogroup. To this end we adapted the analytical pipeline from Grandchamp *et al*. (2023a). Specifically, we first identified the flanking genes of an orthogroup. If an orthogroup did not contain an ORF from a certain line, we extracted the genomic region of the line that lay between the same two flanking genes as the ORFs in the orthogroup. Next, we aligned this region (subject) to the longest ORF in the orthogroup (query) using nucleotide BLAST with a cutoff of 60% query coverage per highest scoring pair (hsp). Next, we identified the highest scoring hit, verified if it is an ORF, translated it into a protein sequence, and mapped it with the longest ORF in the orthogroup using BLASTP (with the same criteria as that we used for transcribed ORFs). If the sequence was successfully mapped with BLASTP, we included it in the orthogroup. We applied this method to every line an ORF from whom was missing in an orthogroup, and then to every orthogroup that had less than seven transcribed ORFs.

### Predicting the ancestral ORF length

To estimate the length of the ancestral ORF for an orthogroup, we used a dated phylogenetic tree of the seven *D. melanogaster* populations (Grandchamp *et al*., 2023b), and the transition matrix estimated using the mutation rate and the intergenic trimer distribution of *D. melanogaster*. We estimate the evolutionary distances in generations from the length of the phlogenetic tree’s branches, assuming that one year has 26 generations (Fernández-Moreno *et al*., 2007). To explain our method better we use the example of a hypothetical orthogroup contains ORF from the Swedish, the Danish and the Spanish populations (leaf nodes), with ORF lengths 50, 50 and 70, respectively. We call the common ancestor of all the three populations “European”, and the a common ancestor of the last two populations, “Scandinavian”. Next, we calculate the number of generations between the European node and each of the three leaf nodes, and use this number to calculate multi-generational transition matrices (*M* ^*n*^). For instance, we start this calculation with the Swedish node (with 254462 generations). Next, we repeat the calculation for the Danish node, but we only count number of generations between the Scandinavian node and the Danish node (161304 generations) because the evolutionary timespan between the European and the Scandinavian nodes (93158 generations) is already calculated for the Swedish node. Finally, we calculate the transition matrix for the evolutionary timespan between the European and the Spanish nodes (254462 generations).

An ancestral ORF can theoretically have any possible length. However, we do not assume a length for the ancestral ORF that is not identical to that of any ORF in the orthogroup. That is because probabilities of extension and truncation are less likely than the probability that ORF length is remains the same, in the evolutionary divergence times between the different *D. melanogaster* populations(Equations 4 – 6). Our consideration minimizes the total number of length changes and excludes unlikely ancestral ORF lengths. Therefore in our hypothetical example, we test ancestral lengths of 50 and 70 and ask which one of them is more likely. First, we calculate the probability that the lengths of the Swedish, the Danish and the Spanish ORFs are 50, 50 and 70, respectively, given that the ancestral length is 50. This would be the product of the corresponding elements of the three transition matrices (as explained in the previous para). We repeat this calculation with now assuming an ancestral ORF length of 70. The ancestral ORF length that gives the largest probability is the most likely ancestor (in this case 50).

We apply the same technique to all the orthogroups that have ORFs with different lengths. Based on the predicted ancestral lengths we calculate the expected number of ORF length changes (truncation, extension, or no change) relative to the ancestor.

### Predicting the expected number of orthogroups with ORFs of different length using Monte-Carlo sampling

With the predicted lengths of the ancestors of every orthogroup, we calculated the probability that they remain the same for the number of generations between ancestral node and extant (leaf) nodes (254462 generations for the hypothetical example used in the previous section). Next, we generated 10^6^ pseudo-random numbers sampled from a uniform distribution with values ranging from 0 to 1, for each of the 758 orthogroups. For each of the 10^6^ iterations, we calculated the total number of random values (out of 758) that exceed the corresponding probability that the ORF length remains the same as the ancestor (Monte-Carlo sampling). This number denotes the expected number of orthogroups that have ORFs with lengths different from that of the corresponding ancestors. In each of the 10^6^ Monte-Carlo samples, we found that the expected number of orthogroups with length difference (median 37), does not exceed the corresponding observed number of orthogroups (173). Hence the observed numbers are greater than expected with a p-value less than 10^*−*6^.

### Analysis of ORF length changes

We analysed length changes in the orthogroups that contained ORFs with different lengths. To this end we first identified the longest ORF for each orthogroup. If there was not one unique longest ORF, we randomly picked one. Next, we compared the longest ORFs to all the other ORFs in the orthogroup using our BLASTP alignments, and classified these ORF pairs into four categories (i) both the ORFs shared both the start and the end positions (ii) both the ORFs shared the same start position but one ORF ended shorter (3’ change) (iii) both the ORFs shared the same end position but one ORF started further than the other (5’ change), and (iv) the shorter ORF was truncated in both sides, sharing neither the start nor the end with the longer ORF.

We analysed categories (ii) and (iii) further. First we analysed the length difference between the short and the longest ORFs (*Δ*). Next, we identified the codons in the longest ORF that overlap the terminal codons of the shorter ORFs. For example, the codon overlapping the stop codon. Conversely, we identified the nucleotide triplet (putative codon) that is located in the 3’UTR, at a distance *Δ* away from the stop codon of the shorter ORF. Likewise, we identified the putative codon in the 5’ UTR of the short ORFs that should positionally align to the start codon of the longest ORF.

### Code and data availability

All source data and analytical codes are freely available. Modeling scripts are available on GitHub:*BharatRaviIyengar/DeNovoEvolution* (specifically Julia scripts ORFlen.jl and analyseTree.jl). Scripts for *Drosophila melanogaster* data analysis are also available on Github: *MarieLebh/ORF_length_evolution*.

